# Defects in the Arabidopsis V-ATPase associated RAVE complex affects endosomal pH and triggers the onset of leaf cell clusters upon TOR inhibition

**DOI:** 10.64898/2026.04.27.721045

**Authors:** Samuel Laurent, Camille Ingargiola, Céline Forzani, Justine Broutin, Isabelle Jéhanno, Claire Perreaux, Gilles Clément, Grégory Mouille, Etienne Delannoy, Jose Caius, Anne-Sophie Leprince, Christian Meyer

**Affiliations:** Institute Jean-Pierre Bourgin for Plant Sciences (IJPB), INRAe, AgroParisTech, Université Paris-Saclay, Centre INRAe Versailles-Saclay, 78000, Versailles, France; Institute of Plant Sciences Paris-Saclay (IPS2), INRAE, CNRS, Université Paris-Saclay, Université Evry, 91405, Orsay, France; UFR 927, Faculté des Sciences et d’Ingénierie, Sorbonne Université, 4 Place Jussieu, 75252 Paris, France

**Author notes:** Corresponding author: Christian Meyer. These authors contributed equally (Samuel Laurent, Camille Ingargiola). Author Contributions: S.L., C.I., C.F., J.B., I.J. and C.P. performed the experiments; G.C. performed the GC–MS metabolites measurements; G.M. performed the cell wall composition analyses; E.D., and J.C. performed the transcriptomic experiment; S.L., C.I., A.-S.L., and C.M. designed the experiments. S.L., C.I., A.-S.L., and C.M. analyzed the data. S.L., and C.M. wrote the manuscript with contributions from all authors. Competing Interest Statement: No competing interests.

**Keywords:** Arabidopsis, V-ATPase, TOR kinase, pH control, cell growth

## Abstract

The TOR kinase is an important and conserved signaling hub in plants, as in other eukaryotes. However, the identification of the TOR pathway components and regulators in plants is still fragmentary. Using a genetic screen based on altered sensitivity to TOR inhibitors, we have selected an Arabidopsis mutant that develops leaf ectopic cell clusters of enlarged cells in a TOR-inhibition dependent manner. We have named this mutant *loki* (Localized growth depending on TOR Kinase Inhibition) and identified the causal mutation in a gene coding for the Arabidopsis homolog of the yeast Rav1 protein. This protein serves as the scaffold for the RAVE complex (Regulator of the ATPase of Vacuolar and Endosomal membranes), which regulates the V-ATPase activity in yeasts and animals. The overall V-ATPase activity is decreased in *loki* mutants and consistently the endosomal pH is increased. However, the vacuolar pH was found to be unaffected by this mutation. Interestingly, the *det3* mutant, which is affected in the C subunit of the V-ATPase, also develops similar cell clusters. Finally, transcriptomic and metabolic analyses revealed that many pathways are affected by both the *loki* and *det3* mutations, including cell wall integrity. This study establishes a new connection between the V-ATPase and the central TOR kinase in plants.

## Introduction

Target of Rapamycin (TOR) is a highly conserved Ser/Thr protein kinase belonging to the Phosphatidyl Inositol Kinase-related kinase (PIKK) family^1^. In yeast and animals, TOR is structuring two complexes, the TOR complex 1 (TORC1) and TOR complex 2 (TORC2). TORC1, composed of the TOR kinase and the two partner proteins RAPTOR and LST8^2,3^ is the better understood of the two. It is a positive regulator of cell division, translation and growth, and a negative regulator of autophagy. TORC2, comprised of TOR, LST8 and RICTOR^3,4^, regulates the actin cytoskeleton as well as cell migration and focal adhesion in mammals, and membrane homeostasis and turgor pressure in yeast cells. In plants, only one TOR complex has been identified^5^. This complex resembles TORC1 but is also involved in plant-specific processes like cell wall assembly, meristem activation or brassinosteroid signaling^6,7^.

The identification of the TOR kinase in yeast was enabled by the discovery of the inhibitor rapamycin and the study of rapamycin-resistant mutants^8^. Rapamycin inhibits TORC1, but not TORC2, in yeast and animals by binding the TOR-interactant FKBP12 thus blocking the kinase activity of the complex^9–11^. Unlike rapamycin, ATP-competitive inhibitors inhibit TOR activity by blocking the ATP binding cleft of the complex and are effective against both complexes^12^. Among those molecules is the inhibitor AZD-8055 (AZD), which was shown to have a high selectivity to TOR compared to other class I phosphatidylinositol 3-kinase (PI3K) isoforms and other members of the PIKK family^13^. In plants, although a homologue of FKBP12 exists, the TOR complex is insensitive to rapamycin^14,15^. However, several ATP-competitive inhibitors have been shown to efficiently and selectively inhibit TOR in plants, including AZD^16^. As knocking-out the TOR kinase is embryo-lethal in Arabidopsis^14^, the study of the complex in plants has benefited from the development of chemical genetic approaches. Pharmacological AZD screens on ethylmethane sulfonate (EMS)-mutagenized populations of Arabidopsis have already led to the characterization of the TOR antagonist AtYAK1^17,18^ and to the identification of AZD-insensitive TOR variants^19^.

Despite those advances, only a small number of regulators upstream of TOR have so far been identified in plants. They include for instance the GTPase Rho-related protein 2 (ROP2) that was shown to activate TOR in response to light or nitrogen^20^, and the cell surface receptor kinase FERONIA (FER) that was found to interact and control TOR activity in a proposed cell wall regulation pathway^21,22^. On the other hand, many upstream regulators of TORC1 activity have been identified in yeast and animals. Notably, the vacuolar H^+^-ATPase (V-ATPase) was shown to be necessary for TORC1 activation and TORC1-mediated nutrient sensing^23,24^. Conversely, TORC1 plays a role in the regulation of V-ATPase activity^25^. In animals, inhibition of TORC1, by either amino acid (AA) starvation or inhibitor treatment, leads to an increase in V-ATPase assembly at the lysosome which results in an increased V-ATPase activity and subsequent acidification at this compartment^25^. As in mammals, yeast TORC1 is able to regulate the assembly state and activity of the complex in a nutrient-dependent manner via the downstream effector Sch9^26–28^.

V-ATPases are highly conserved transmembrane proton pumps responsible for the acidification of endomembrane compartments of eukaryotic cells. They are comprised of two domains, a transmembrane domain that serves as a proton channel: V_0_, and a cytosolic domain responsible for ATP hydrolysis: V_1_. Each domain contains several subunits ranging from VHA-a to VHA-e (V_0_ domain) and from VHA-A to VHA-H (V_1_ domain)^29^ with some subunits existing as multiple isoforms that dictate spatial partitioning of V-ATPases and V-ATPase function. In yeast, VHA-a has two isoforms: Vph1p is vacuole-specific and Stv1p that localizes to the Golgi^30,31^. As a result, the disruption of a single one of those isoforms leads to specific phenotypes^32,33^. Similarly, in Arabidopsis, V-ATPases localize to various compartments of the endomembrane system^34^. This compartment specificity of V-ATPase activity is essential for a number of biological processes as evidenced by the published studies on various V-ATPase mutant lines with distinct phenotypes^35^. Notably, some V-ATPase mutants are specifically affected at the Trans Golgi Network/Early Endosome (TGN/EE). For instance, expression of inducible RNAi targeting VHA-a1, the TGN specific isoform of VHA-a, leads to strong inhibition of hypocotyl elongation and cell expansion, while the tonoplast specific *vha-a2xvha-a3* double mutants do not display severe cell expansion defects^36–38^.

Aiming to identify molecular components of the TOR signaling pathway, we undertook a pharmacological screen to isolate new mutants hypersensitive to the inhibition of TOR by AZD. We identified the *loki* mutant, a novel mutant displaying hypersensitivity to AZD as well as an ectopic cell cluster phenotype. We found that the causal mutation lies in the sequence of a gene homologous to the yeast V-ATPase regulator Rav1^39^ pointing toward a link between TOR and V-ATPases in Arabidopsis. We show a strong phenocopy between this new mutant and the TGN-affected V-ATPase mutant *det3*^40,41^, associated with similar transcriptomic responses to AZD. This study aims to propose a first investigation and characterization of the *loki* mutant and its ectopic cell cluster phenotype. We also propose hypotheses in the role of LOKI in V-ATPase regulation.

## Results

### An AZD screen identifies an Arabidopsis homologue of the yeast protein Rav1

In order to identify new actors of the TOR pathway in Arabidopsis, a screen was undertaken on a population of double-haploids fixed for EMS mutations^42^ to isolate mutations inducing hypo- or hyper-sensitivity to the pan-PI3K inhibitor LY294002 (LY)^16^ and the TOR-specific inhibitor AZD8055 (AZD). Among the hyper-sensitive mutants identified in this screen, one line, HD305, stood out as it displayed ectopic cell clusters growing on aerial organs upon TOR inhibition (Fig. 1A). These clusters were never detected on roots and the HD305 line was the only one to show this particular phenotype. The HD305 line was backcrossed in a wild-type (WT) Col-0 background and descendants displaying the mutant phenotype were sequenced in bulk. The MutDetect pipeline^43^ was used to identify sequence polymorphisms. To map the causal mutation of the HD305 line, we used the EMS-induced sequence mutations to detect regions in the Arabidopsis genome highly enriched in homozygous polymorphisms and putatively associated with the HD305 mutation due to linkage disequilibrium. The bottom of chromosome 2 displayed a high proportion of homozygous sequence polymorphisms (Fig. 1B) and 10 homozygous null mutations were identified in this region.

**Figure 1.**
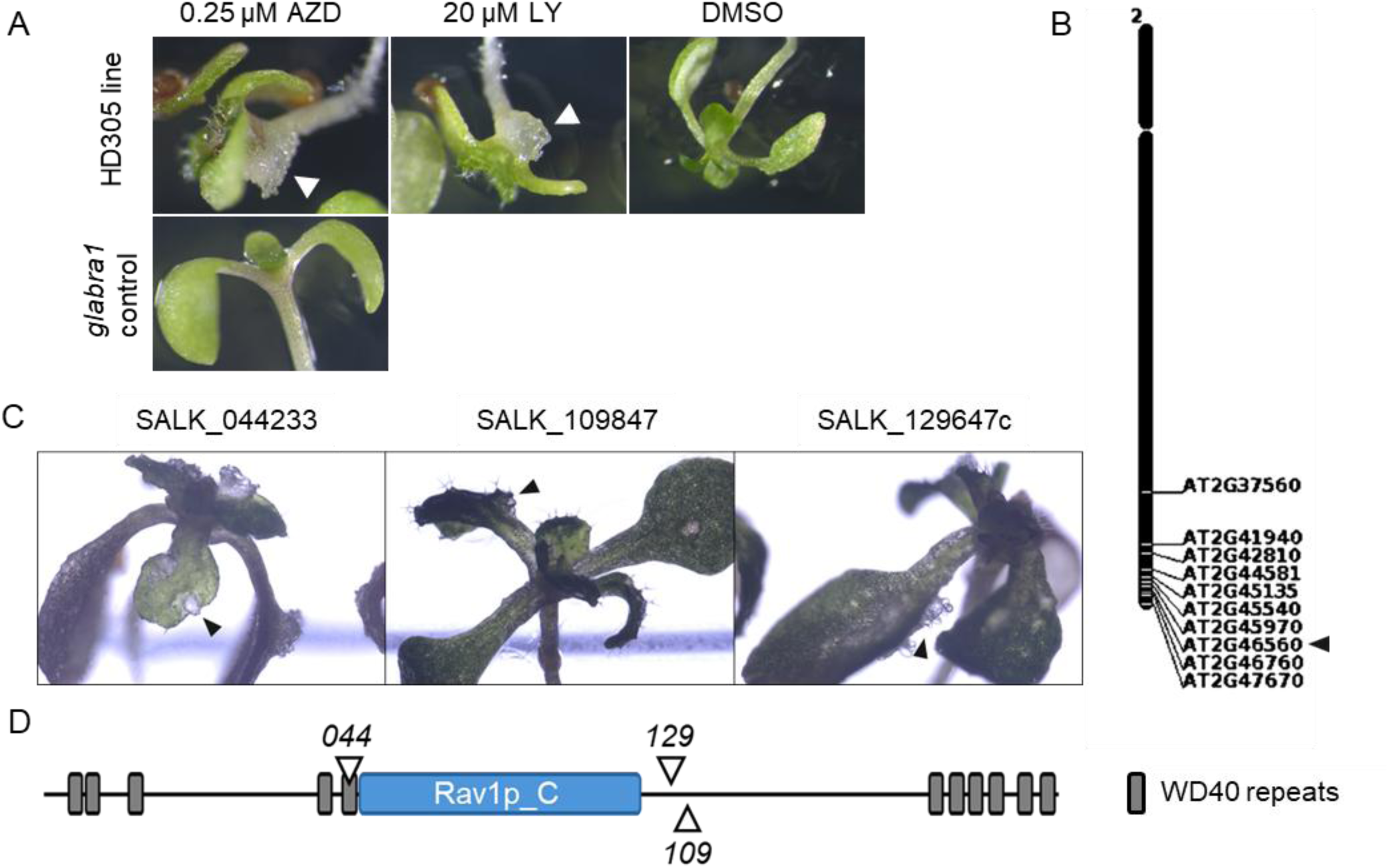
Identification of the *loki* mutant that displays an ectopic cell cluster phenotype upon TOR inhibition. (A) Phenotype of the HD305 double haploid line on AZD8055 (AZD, 0.25μM), LY292004 (20μM) or DMSO. This line is in the *glabra1* background. Ectopic cell clusters are indicated by arrowheads when present. (B) The bottom of chromosome 2 was enriched in homozygous mutations in a progeny of a cross between *glabra1* and HD305 lines when plants carrying ectopic clusters were selected and sequenced in bulk. (C) T-DNA insertion mutants corresponding to the *AT2G46560* gene exhibited the same cell cluster phenotype (indicated by arrowheads) when treated with 0.25µM AZD. (D) The LOKI protein encoded by this gene carries WD40 domains and a central domain homologous to the yeast Rav1 domain (Rav1p_C, pfam12234). The T-DNA insertions of SALK_044233, SALK_109847 and SALK_129764c, named *044*, *109* and *129* respectively, are shown relative to the WD40 and Rav1 domains of LOKI. The LOKI domain information was retrieved from InterPro (https://www.ebi.ac.uk/interpro/).

We selected the three most promising candidates among these genes by their links to known regulatory elements of the TOR kinase in other organisms. T-DNA insertion mutants corresponding to these candidate genes were obtained and homozygous mutants were tested for the formation of cell clumps upon TOR inhibition. Three independent T-DNA mutants affected in the *At2g46560* gene developed the same phenotype as the HD305 mutant on AZD (Fig. 1C). The causal mutation in the HD305 line was then identified as a stop mutation in the gene *At2g46560* (G→ A mutation changing the Trp 552 residue into a stop codon). The T-DNA insertion sites of the three *at2g46560* alleles are shown in Fig. 1D. The absence of the *At2g46560* transcript in each line was confirmed by RT-PCR with primers spanning the length of the coding sequence (Fig. S1A).

The *At2g45560* locus encodes a long 2513 amino acid (AA) protein with a molecular weight (MW) of 275kDA. We named this hitherto unknown protein LOKI for ‘LOcalized growth depending on TOR Kinase Inhibition’. LOKI carries three WD40 repeats-containing domains, two at the N-terminus and one at the C-terminus (Fig. 1D, Fig. S1C). The central part of the LOKI protein sequence is homologous (26% identity) to the Rav1 domain of the yeast Rav1 protein (Fig. S1C, Fig. S2). Rav1 is a smaller protein (1357 AA) that also contains two WD40 domains at the N-terminus and the α solenoid domain Rav1, but only a short, disordered tail at the C-terminus. Rav1 homologues also exist in animals (Rabconnectin-3a, DMXL1/2) with the mammalian proteins being much larger in size (3027 AA and 3036 AA for the human proteins HsDMXL1 and HsDMXL2 respectively) than Rav1 or LOKI, and containing an enlarged Rav1 domain in the central region^44^. Phylogenetically, LOKI was found to be closer to HsDMXL1 and HsDMXL2 than to the yeast Rav1 (Fig. S1B) with 32% identity at the Rav1 domains (Fig. S1C, Fig. S2). In yeast, Rav1 is a member of the RAVE (Regulator of the ATPase of Vacuolar and Endosomal membranes) complex together with the Rav2 and Skp1 proteins^45^. This complex, also characterized in animals, is involved in regulating the reversible assembly of V-ATPase V_1_ and V_0_ domains^39^.

### *loki* mutation results in ectopic cell cluster phenotype upon TOR-inhibition

To understand the link between V-ATPase regulation and the observed cell clusters phenotype, we first characterized these ectopic growths. After two weeks, about half of AZD-grown *loki* seedlings displayed these transparent cell mass at the leaf surface (Fig.S3A). This rate of apparition was about the same for all three *loki* knockout lines, thereafter referred to as *loki044*, *loki109* and *loki129*. Those clumps were never observed on the root system. To confirm the link between TOR inhibition and the onset of the cell cluster phenotype, the *loki109* mutant was crossed with the *lst8-1* mutant^46^, affected in TOR activity. The *loki109xlst8-1* double mutants displayed the ectopic cell cluster phenotype even in the absence of any chemical TOR inhibitor (Fig. S3B). The *loki* mutants were also grown on another specific TOR inhibitor, Torin2. The cell cluster phenotype was again observed (Fig. S3C).

To visualize the anatomy and morphology of those *loki* cell clusters, calcofluor stained cluster-bearing leaves were examined under a confocal microscope using a long-distance objective to maintain the integrity of the cluster. This showed that those clusters were composed of expanded cells swelling from beneath the leaf epidermis (Fig. 2A). The fully expanded cells were visibly devoid of chlorophyll (Fig. S4A), which was reminiscent of the structures observed in heat-induced mesophyll cell expansion^47^. It has been shown that those cell expansion events following heat stress were dependent on programmed cell death. Interestingly, assessing cell viability by dual staining with fluorescein diacetate (FDA, which stains live cells) and propidium iodide (PI, which stains dead cells) showed that small clusters were comprised of live cells with enzymatic activity and intact membranes while larger, older clusters contained some live cells but also some dead cells in which PI stained the nucleus (Fig. 2D). Those *loki* cell clusters were primarily observed on the path of the vasculature (Fig. 2B, Fig. S4B). Closer examination of DirectRed stained leaves revealed a close association between the expanded cells and vascular tissues (Fig. 2C), while no defects in vascular tissues organization were observed in *loki* leaves from seedlings grown in standard conditions (Fig. S4C). This correlates with Single Cell data (https://bioit3.irc.ugent.be/plant-sc-atlas/)^48^ that predicts *LOKI* to be most expressed in the leaves’ guard cells, xylem and phloem parenchyma (Fig. S4D).

**Figure 2.**
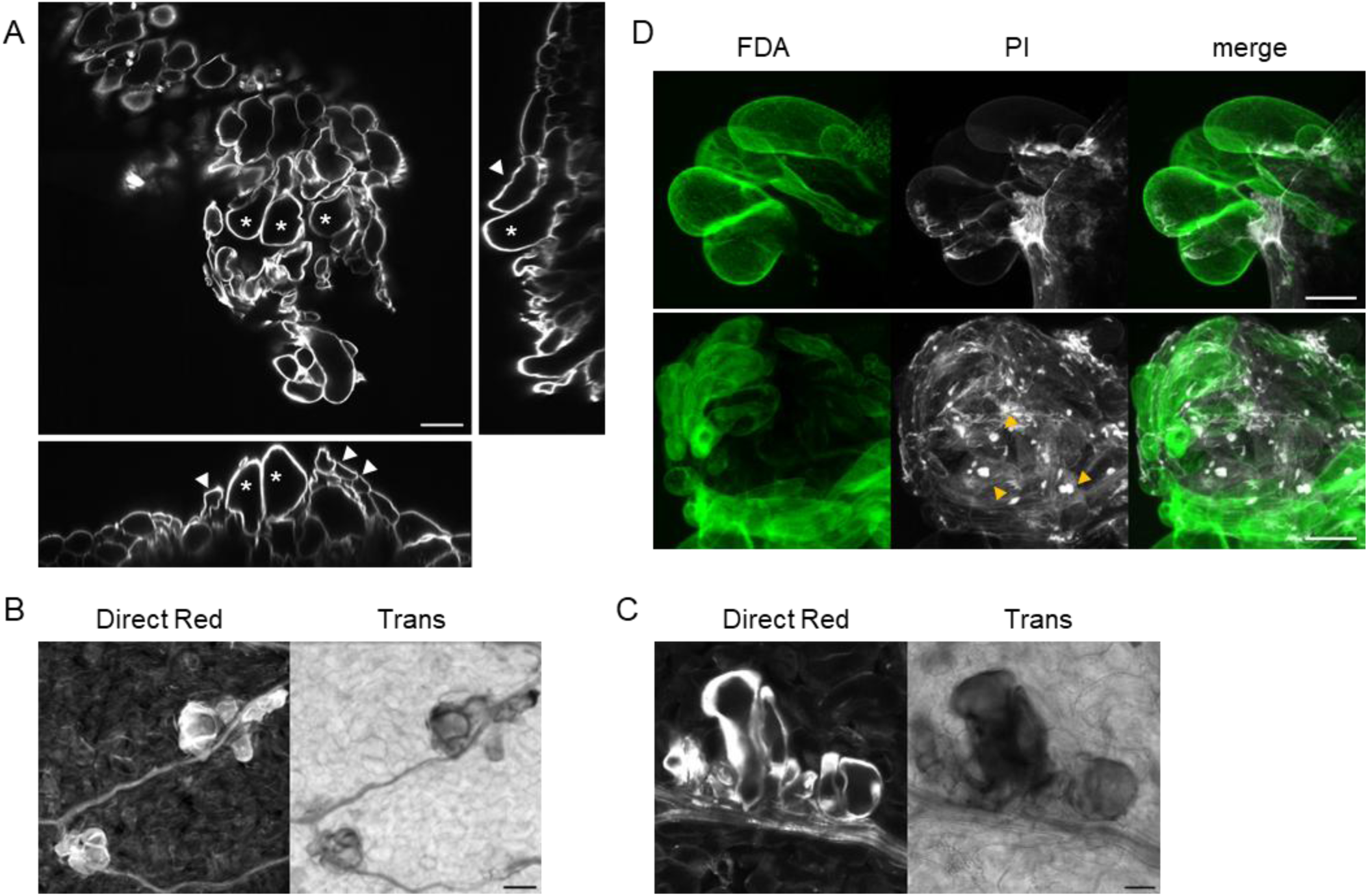
Morphological and cytological characterization of the ectopic cell clusters. (A) Section of a calcofluor-stained cluster of expanded cells from a leaf of a *loki044* mutant with reconstructed orthogonal slices (asterisks indicate expanded cells, arrowhead indicate epidermal cells), scale bar = 100µm. (B, C) Confocal and associated transmitted light (Trans) images of cell clusters stained with Direct Red. Images were acquired on two different plants, scale bars = 100µm (B) and 50µm (C). (D) Representative confocal images of expanded cells labeled with fluorescein diacetate (FDA, green) and propidium iodide (PI, white) from different types of clusters. Arrowheads indicate propidium-iodide-stained nuclei, scale bar = 100µm.

### *loki* mutants are hypersensitive to TOR inhibition

Beyond the ectopic cell cluster phenotype, we tested whether *loki* mutants showed any other specific phenotype upon TOR inhibition. Since TOR was shown to be necessary for hypocotyl elongation in the dark^7^, we measured hypocotyl lengths of dark-grown *loki* mutants on 0.25µM of AZD. DMSO-grown *loki* mutants had only slightly decreased hypocotyl lengths compared to the WT. But on AZD, *loki* mutants showed a 60% reduction in hypocotyl elongation while the length of WT plants hypocotyls was only reduced by 21% (Fig. 3A). This hypersensitivity to TOR inhibition was also visible in the projected rosette areas (PRA) of seedlings (Fig. 3B). Interestingly, root length of *loki* mutants was less sensitive to AZD treatment (Fig. S5). To assess whether this hypersensitivity to TOR inhibition was due to an altered basal TOR signaling in *loki*, we measured TOR activity after induction by sucrose or inhibition by AZD. Phosphorylation levels of the TOR downstream target Ribosomal Protein S6 (RPS6) in the *loki* mutant showed no significant difference compared to the WT in their response to TOR induction and inhibition (Fig. 3C).

**Figure 3.**
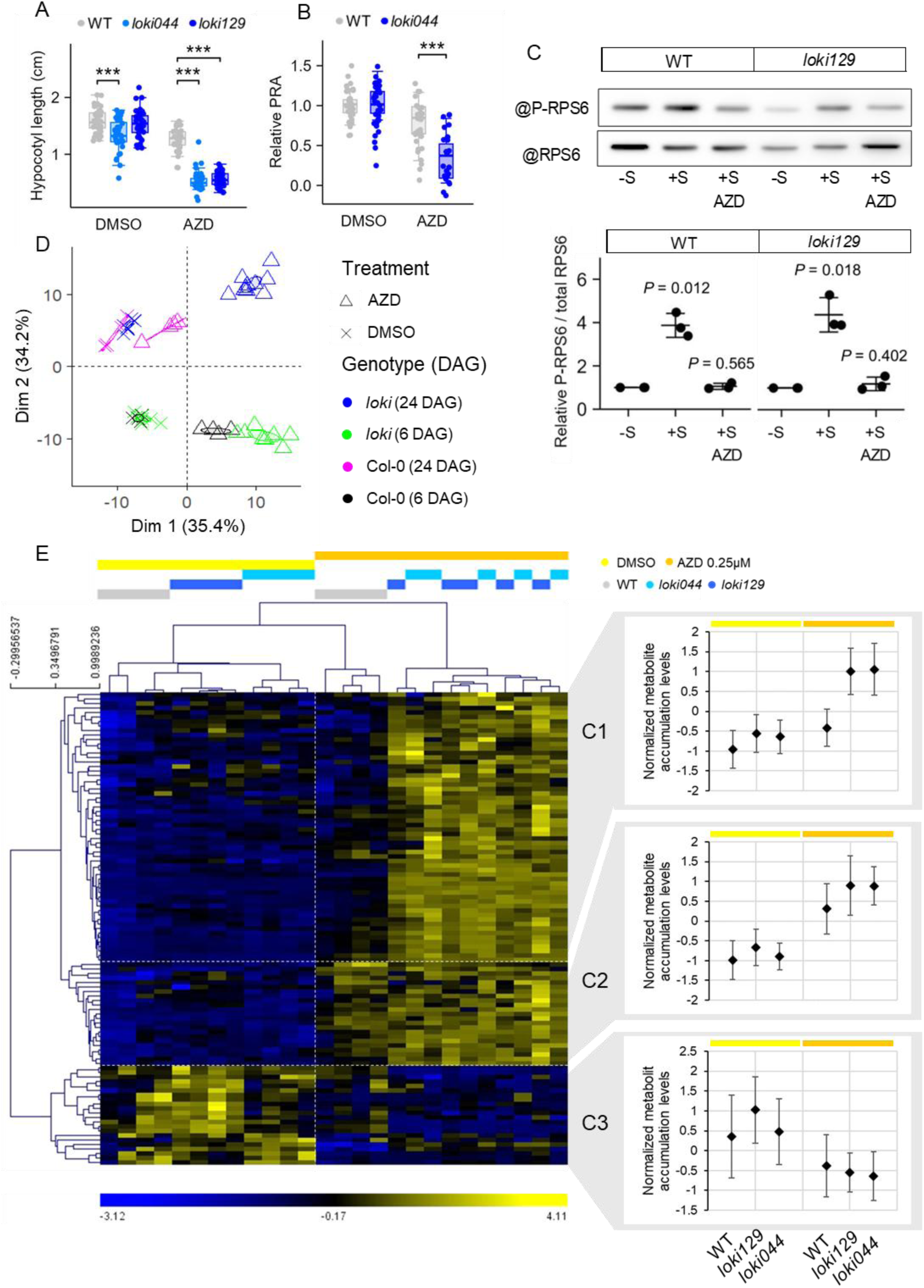
The *loki* mutants show growth defects and metabolic changes upon TOR inhibition. (A) Hypocotyl lengths measured on dark-grown 14-day-old seedlings in the presence of 0.25µM AZD or equivalent amounts of DMSO for control. *n* = 35-40. (B) Relative Projected Rosette Area (PRA) of 12-day-old seedlings grown in the presence of 0.5µM AZD or equivalent amounts of DMSO. *n* = 25-36. (A, B) Asterisks indicate statistical significance, \*\*\**P* < 0.001. *P*-values were calculated using a Dunn statistical test with BH correction. (C) Phosphorylation levels of RPS6 reflecting TOR activity. Representative membrane revelation (top) and quantification (bottom) from three independent experiments. –S: sucrose starvation conditions, +S: sucrose induction conditions, +S AZD: sucrose induction medium supplemented with 1µM AZD. *P*-values were calculated using a Student’s t-test with BH correction. (D) PCA of metabolite data from seedlings grown for 6 days or 24 days on 0.25µM AZD (triangles) or equivalent amount of DMSO (crosses). (E) Heatmap generated from a hierarchical clustering analysis of changes in metabolite abundance between genotypes and treatments at 24 days after germination (DAG). Significantly over- or under-accumulated metabolites were determined with a two-factor Mark-Sillings test (alpha = 0.05). The hierarchical tree was drawn from significant metabolites (Distance metric: Pearson correlation, Linkage method: average linkage clustering). The scale bar (bottom) reflects the fold changes of metabolite accumulation. Based on accumulation patterns, AZD-responsive metabolites were divided into three clusters. The plots display the normalized accumulation levels within each cluster.

The inhibition of the TOR pathway leads to numerous changes in the metabolome of a plant. Amino acids, lipids and TCA cycle intermediates are notably affected, as well as storage components such as TAGs^46,49–51,18^. To determine whether the hypersensitivity at the whole plant level was associated with altered metabolic responses to TOR inhibition, we performed metabolite profiling on seedlings grown for either 6 days (before the onset of the cluster phenotype) or 24 days (past the onset of the cluster phenotype). We compared the responses in the metabolite profiles of WT plants and the *loki* mutants grown under control condition or with 0.25µM of AZD. For a visual inspection of the extent of those metabolomic changes under AZD treatment, we subjected the metabolite profile data to PCA (Fig. 3D). On the PCA score scatter plot, the samples were clustered according to both age and treatment. We observed a marked genotype-dependent separation only between 24-days old AZD-treated plants.

To further dissect the metabolic response of *loki* mutants on AZD, we categorized the treatment-responsive metabolites at 24 days into three clusters (Fig. 3E). Cluster 1 (C1) includes metabolites known to increase in response to AZD in the WT but that over-accumulate in *loki* like AAs, especially branched-chain AAs (BCAAs), and polyamines like putrescine and its precursor, agmatine (Fig. 3E, Fig. S6A). Cluster 1 also contains numerous soluble sugars, intermediates of starch biosynthesis and pectin precursors like galacturonate, whose accumulation upon TOR inhibition was attributed to hindered cell wall assembly upon TOR inhibition^50^. Finally, metabolites that accumulate in *loki* but not in Col-0 in response to AZD are also found in Cluster 1, this is the case of numerous stress metabolites, notably ROS scavengers (Fig. 3E, Table S2). Cluster 2 (C2) comprises of metabolites that over-accumulate in response to AZD in a similar manner in both *loki* and Col-0. It includes some AAs (Tyr, Gly, Trp, Arg, Asp, Glu, Asp, homoserine) and a few sugars like xylose and galactose (Fig. 3E, Fig. S6B). Cluster 3 (C3) consists of metabolites that are depleted in response to AZD. It includes metabolites like raffinose and myo-inositol with similarly decreased levels in *loki* and Col-0 as previously described^46^. Although TCA intermediates have been shown to accumulate in Arabidopsis in response to AZD^49,50^, only citrate is found in Cluster 2 (Fig. 3E, Table S2) while fumarate and succinate are found in this third cluster (Fig. 3E, Fig. S6C, Table S2). A few metabolites are depleted in Col-0 in response to AZD but not in *loki* like stigmasterol. The only metabolites found to be more depleted in *loki* than in the WT were the fatty acids linoleic acid and linolenic acid (Fig. 3E, Fig. S6C).

Consistently with observations based on the PCA scatter plot, clustering samples from 6-day-old seedlings based on metabolite profile did not allow for discrimination between *loki* and Col-0 (Fig. S7A). Nonetheless, hierarchical analysis highlighted a cluster of metabolites with increased levels in *loki* compared to the WT on AZD (C1’; Fig. S7A). Among the metabolites found in this Cluster 1’, 60% are also found in Cluster 1 at 24 days (Fig. S7B). These shared metabolites include the three BCAAs (Leu, Ile, Val), agmatine and putrescine, and several glucosinolates but only a few sugars. Other oxidative stress responsive metabolites are also shared between the two sub-clusters (tocopherols, ascorbate). Overall, these results suggest that the *loki* mutation by itself does not affect the global metabolism at the whole plant level but does result in starker responses to TOR inhibition.

### *loki* loss-of-function phenocopies *det3* knockdown

Because the *LOKI* protein is homolog to Rav1, we then investigated whether these phenotypes could be linked to changes in V-ATPase activity. In yeast, *rav1*Δ mutants have structural and functional V-ATPase defects which result in “partial” Vma- phenotypes, i.e. similar but less intense phenotypic traits as mutants lacking V-ATPase subunits^45,52^. To test whether the cell cluster phenotype is a result of V-ATPase dysfunction, we sowed various V-ATPase mutants on AZD-containing medium. Interestingly, the *det3* mutant (carrying a weak allele for the VHA-C subunit^40^), displayed similar cell clusters on AZD, while the double mutant *vha-a2xvha-a3*^38^ did not (Fig. 4A, Fig. S8A).

**Figure 4.**
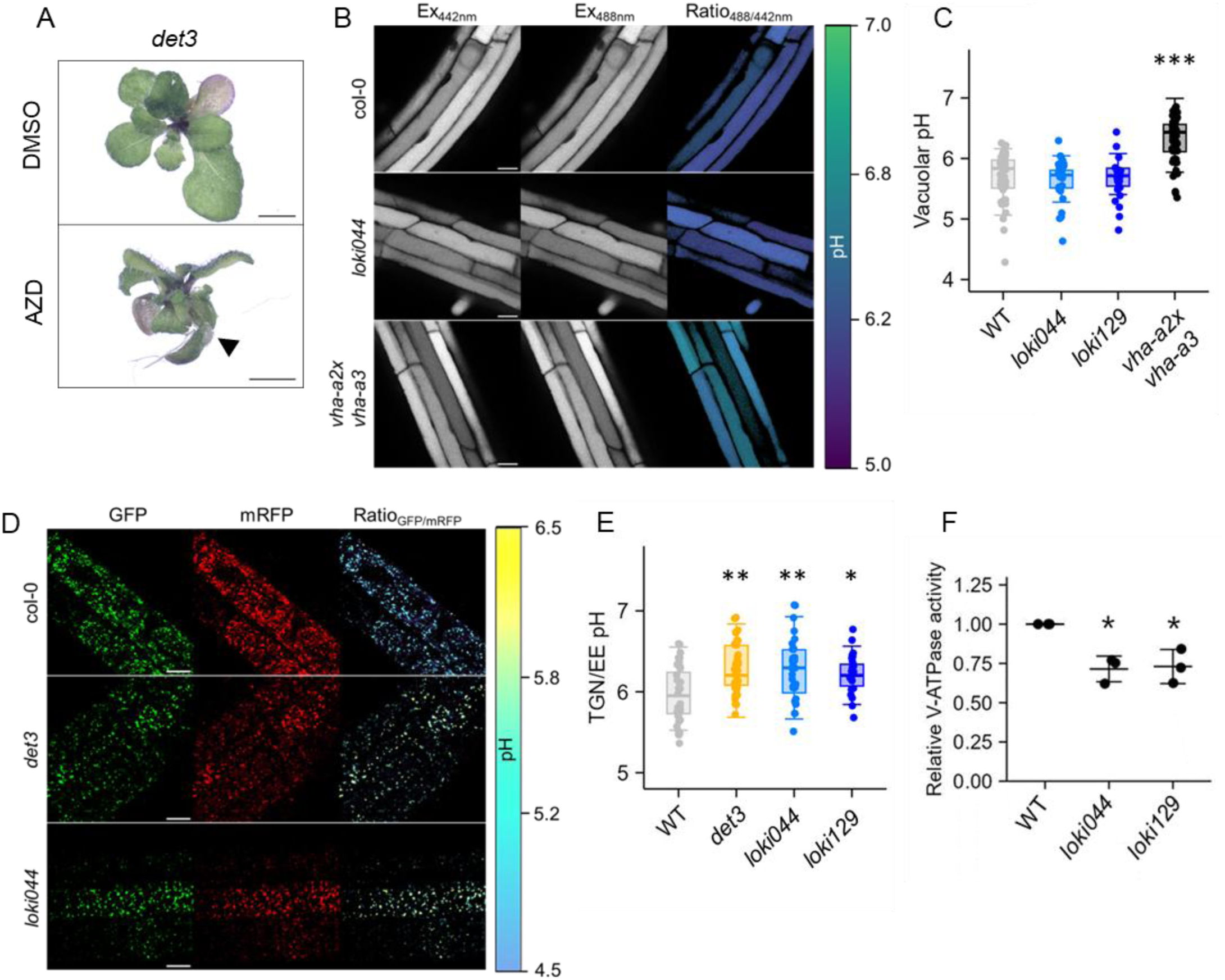
*loki* and *det3* display similar defects in endosomal pH and ectopic cell clusters on AZD. (A) Representative image of *det3* seedlings grown on 0.5µM AZD or DMSO. Ectopic cell clusters are indicated by arrowheads when present, scale bar = 2cm. (B) Emission intensities of epidermal root cell vacuoles loaded with BCECF,AM. The ratio image of *vha-a2xvha-a3* shows an increased vacuolar pH, scale bar = 100μm. (C) Vacuolar pH measured from BCECF,AM stained root epidermal cells (B), *n* ≥ 27. (D) Emission intensities of TGN/EE vesicles from epidermal root cells expressing the SYP61-pHusion construct. The ratio images of *loki044* and *det3* show an increased TGN/EE pH, scale bar = 100µm. (E) TGN/EE pH measured from root epidermal cells expressing SYP61-pHusion (D), *n* ≥ 26. (F) Nitrate- and Concanamycin A-inhibited V-ATPase activity measured from microsomal membranes of 7-day-old seedlings. Wild-type activity was set to 1. (C, E, F) Asterisks indicate statistical significance, \**P* < 0.05, \*\**P* < 0.01, \*\*\**P* < 0.001. *P*-values were calculated using a Dunn statistical test with BH correction.

To investigate if the LOKI protein might influence vacuolar pH through its involvement in V-ATPase regulation, we measured vacuolar pH in the *loki* mutants using the ratiometric pH indicator 2,7-bis-(2-carboxyethyl)-5-(and-6)-carboxyfluorescein, acetoxymethyl ester (BCECF,AM). While vacuoles in *vha-a2xvha-a3* double mutants had the expected increased pH^38^, pH in *loki* vacuoles was indistinguishable from WT pH (Fig. 4B, C) suggesting that V-ATPase activity at the tonoplast in *loki* mutants is either unaffected or compensated for by other mechanisms.

*det3* knockdowns have been shown to display phenotypes typically associated with V-ATPase defects at the TGN/EE^40,37^. Because the *det3* response to AZD was similar to that seen in *loki*, we investigated whether the pH of the TGN/EE was also affected in *loki* mutants. We crossed the *loki044* and *loki129* mutants with a reporter line expressing the ratiometric pH sensor pHusion^53^ fused to the TGN/EE marker SYNTAXIN OF PLANTS61 (SYP61)^41^. We measured a pH of 6.15 in both *loki* mutants, a value similar to the one measured in *det3*. By contrast, TGN pH in the WT was measured at 5.81 (Fig. 4D, E). We confirmed those pH defects as due to V-ATPase dysfunction with an activity assay. Total V-ATPase activity was measured as Concanamycin A and NO_3-_-sensitive ATP hydrolysis^38^ in microsomal membranes from whole WT and *loki* seedlings. Compared to the wild-type, V-ATPase activity in both *loki* mutants was reduced by about 25% (Fig. 4F). Consistent with our observations, it was shown that the double mutant *vha-a2xvha-a3*, in which only the TGN-specific VHA-a1-containing V-ATPase complexes are active, displays a 80-85% reduction in V-ATPase activity compared to the WT^38,54^. This further suggests that only V-ATPases at the TGN/EE are affected by the *loki* mutation.

In animal cells, inhibition of TOR leads to an increase in V-ATPase activity that is dependent on RAVE-mediated reassembly^25^. To assess whether disruption of TOR pathways in Arabidopsis has similar effects, we measured vacuolar and TGN pH of AZD-grown seedlings. We found that chronic TOR inhibition had no effect on vacuolar pH in either the WT or *loki* mutants, nor on TGN pH of the WT (Fig. S8B). However, AZD treatment led to a worsening of pH alkalinization at the TGN in *loki* (Fig. S8C).

### Transcriptome shows similar responses to AZD in *loki* and *det3*

To gain further insight into the molecular processes involved in the cluster onset and AZD response in *loki* and *det3*, we looked at the transcriptomic profiles of 18-day-old WT, *det3* and *loki* mutants grown under control condition or with 0.25µM of AZD (Fig. S9A, B). The number of differentially expressed genes (DEGs) on AZD compared to DMSO was much higher in both *loki* mutants than in the WT (Fig. S9A, B), reinforcing previous observations of their hypersensitivity to TOR-inhibition. As no significant difference was found in the transcriptomic responses to AZD between *loki044* and *loki129*, they were grouped for further analyses. We found a significant correlation (Pearson test *P*-value < 2.2 10^-16^) between the transcriptomic response to AZD of both *loki* and *det3* (Fig. 5A). Among the differentially expressed genes in response to AZD in *loki* compared to Col-0, 56.8% of downregulated genes (Fig. 5C) and 43.6% of upregulated genes (Fig. 5B) were also found in AZD-grown *det3*. Moreover, 9.9% of downregulated genes and 30.1% of upregulated genes in AZD-grown *loki* were found in both DMSO- and AZD-grown *det3*. GO term analysis (https://pantherdb.org/) of those shared AZD-responsive DEGs showed an enrichment of various biotic and abiotic stress response pathways, including response to hypoxia, response to wounding, response to oxidative stress, response to water deprivation and detoxification (Fig. 5D). This similarity in transcriptomic responses between *loki* and *det3* suggests a similar mechanism underlying the ectopic cell cluster phenotype.

**Figure 5.**
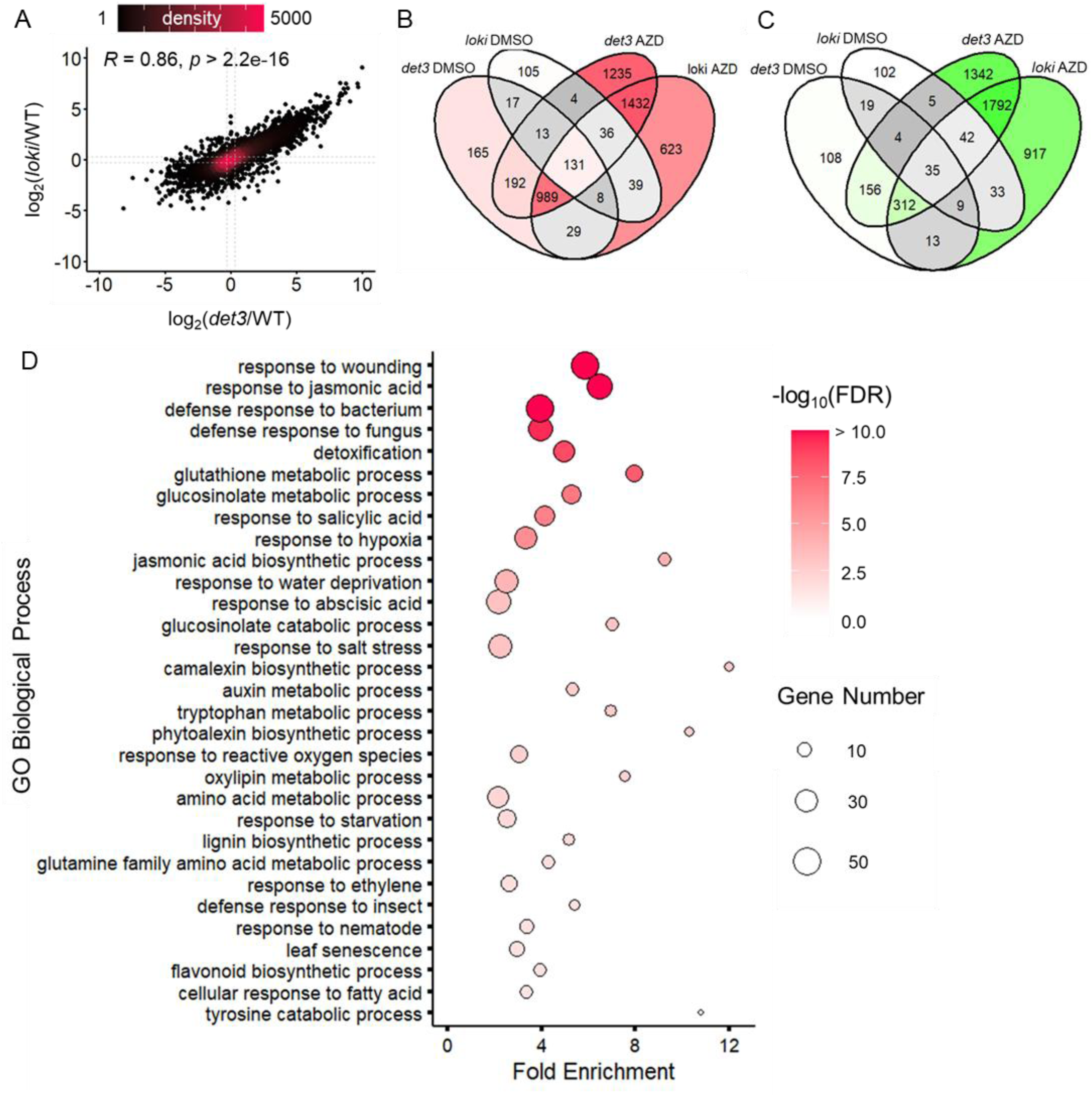
Transcriptomic analysis of *loki* and *det3* responses to AZD show hypersensitive and overlapping responses. Transcriptomic analyses from seedlings grown for 18 days on 0.25µM AZD or equivalent amounts of DMSO for control. (A) Correlation analysis of AZD transcriptomic response in *det3* and *loki* compared to the WT response. Each gene is represented as a dot. The x axis represents difference in gene expression fold change between AZD-treated *det3* and AZD-treated Col-0. The y axis represents difference in gene expression fold change between AZD-treated *loki* and AZD-treated Col-0. Color represents dot density. *R* indicates the calculated Spearman’s correlation coefficient. (B, C) Venn diagrams of genes significantly upregulated (B, log_2_(FC) > 0.3, *P*-value < 0.05) and downregulated (C, log_2_(FC) < -0.3, *P*-value < 0.05) in *det3* and *loki* compared to Col-0 under the same treatment conditions. Color intensity represents gene counts. (D) Gene Ontology analysis of the 1432 genes upregulated on AZD in both *loki* and *det3* compared to Col-0 (B). Bubble size represents the number of genes, color indicates significance (False Discovery Rate).

We also performed transcriptomic analyses using laser-assisted microdissection and capture of cell clusters to further dissect the genes and molecular mechanisms involved in the onset of cell clusters. Captured clusters were obtained for *loki044* and *loki129* mutants. As a control, we used microdissected leaf samples of similar sizes that did not display any visible cell clusters, harvested from the same AZD-treated mutants. 170 and 167 genes were found to be up-regulated in, respectively, *loki044* and *loki129* cell clusters (log_2_(FC) > 3, Fig. S10A). Among these induced genes, 123 were common to the two mutant lines which shows a good reproducibility of this analysis. A meta-analysis (http://bioinfo.sibs.ac.cn/plant-regulomics/) of these results show that induced genes are rather similar to genes induced during callus formation (56 genes out of 123), wound response or drought stress.

### TOR inhibition leads to cell wall defects in *loki*

Grouping *loki* AZD-responsive DEGs by cellular compartment showed an enrichment of genes associated with the cell wall and the endomembrane system (Fig. S11). The *det3* mutation is known to result in changes of cell wall composition^55,41^. It was shown that the *det3* mutation causes a global decrease in intracellular trafficking speed, including a reduction of cellulose biosynthesis cesA complexes secretion which results in a decreased cellulose content^41^. This leads to a reduced cell wall stiffness in *det3* mutants and a reduced constraint exerted by the epidermal layer on the inner tissues. In the *clv3xdet3* double mutant, these defects combined with an increased cell division activity causes the stem to crack^56^.

Given the similarities between *loki* and *det3*, we proposed the hypothesis that the ectopic cell clusters seen upon AZD treatment could result from similar mechanical defects in combination with changes induced by TOR inhibition. Measuring cellulose content in *loki* mutants showed a slight decrease compared to the WT that was worsened in AZD-treated seedlings (Fig. 6A). Consistently, we found that *loki* mutants were hypersensitive to the inhibition of cellulose biosynthesis by isoxaben (Fig. 6B). Scanning electron microscopy showed collapsed epidermis cells in hypocotyls of AZD-treated *loki* plants (Fig. 6D), likely stemming from a loss of cell wall rigidity similar to the one described in *det3*^56^. To confirm that the cellulose content reduction in *loki* mutants results from the same processes as in *det3*, we quantified the uptake and intracellular kinetics of the fluorescent styryl dye FM4-64, routinely used as endocytic tracer in plants, in root epidermal cells. In *det3*, the delayed intracellular trafficking has been shown to result in an accumulation of FM4-64 in the endosomal compartment leading to an increase in the relative intracellular fluorescent signal of the dye, only at late timepoints^41^. No significant differences between WT and *loki* mutants were observed at early time points indicating a normal rate of endocytosis in the mutants (Fig. 6F, 6G). However, at later timepoints, FM4-64 signal was clearly found at the tonoplast in WT cells but still mostly labelling endosomes in *loki* cells, suggesting a delayed endocytic trafficking similar to what is observed in *det3* mutants (Fig. 6F, G).

**Figure 6.**
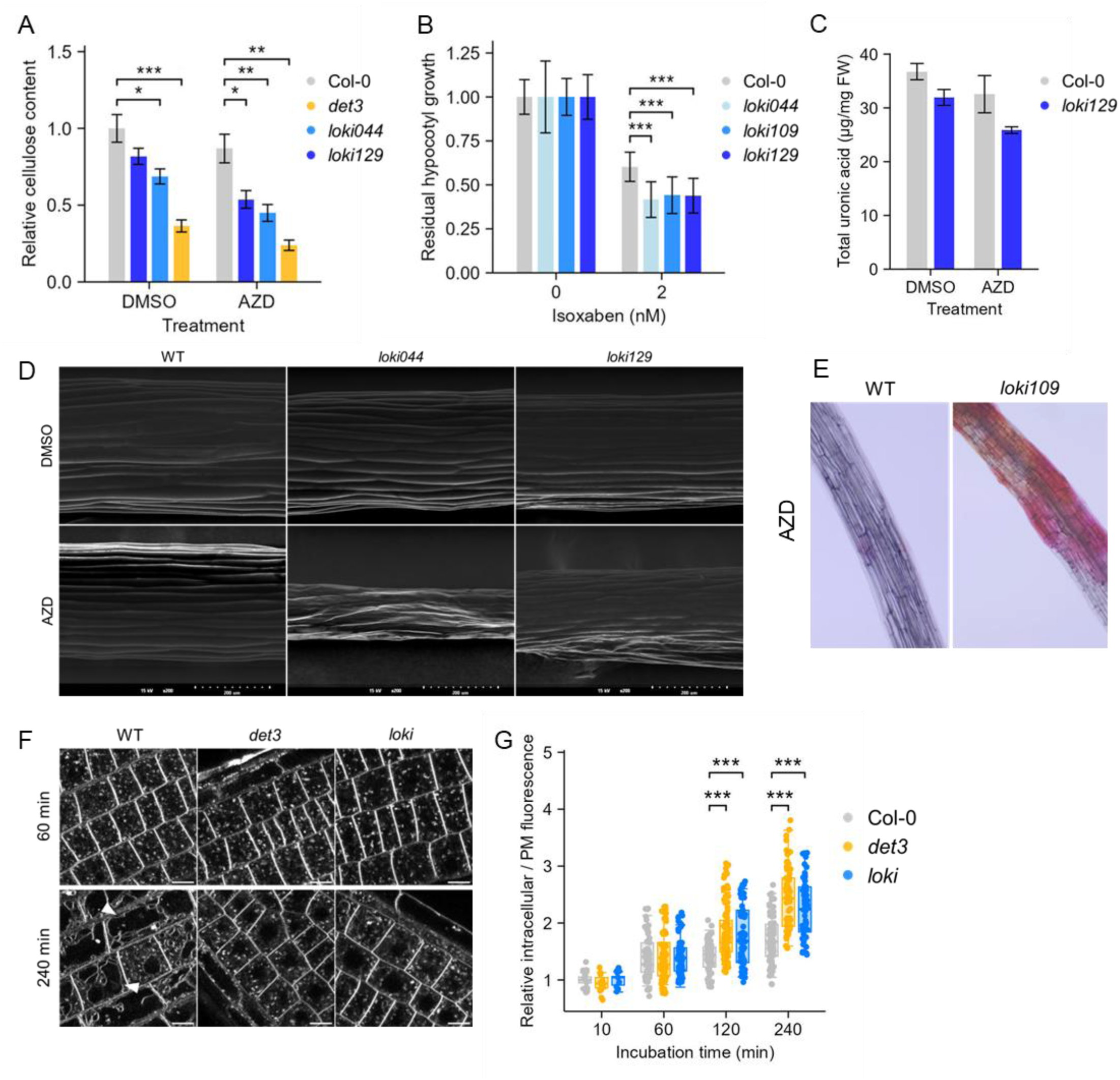
*det3* and *loki* mutations affect cell wall composition and trafficking. (A) Relative cell wall cellulose content measured from 14-day-old hypocotyls of Col-0, *loki044*, *loki129* and *det3* grown on either DMSO or 0.25µM AZD. Measurements were done on *n* = 4-8 samples per condition, each sample consisting of 52-153 hypocotyls. Error bars represent SEM. (B) Residual hypocotyl lengths measured on seedlings grown for 14 days on 2nM of isoxaben or equivalent volume of methanol for control (0nM). Error bars represent SD across three repetitions, *n* = 17-33. (A, B) Asterisks indicate statistical significance between mutant lines and the WT, \**P* < 0.05, \*\*\**P* < 0.001. *P*-values were calculated using a Student’s t-test with BH correction. (C) Total uronic acid content of cell wall material from Col-0 and *loki129* hypocotyls. Measurements were done on two samples per condition and genotype, each sample consisting of 36-107 hypocotyls. Error bars represent SEM. (D) Representative scanning electron microscopy images of 3-day-old hypocotyls of Col-0, *loki044* and *loki129* grown on DMSO or 0.25µM AZD. (E) Ruthenium Red staining of 14-day-old hypocotyls of Col-0 and *loki109* grown on 0.25µM AZD. (F, G) FM4-64 uptake at 10min, 1h, 2h and 4h quantified as intracellular fluorescence intensity normalized to the fluorescence intensity at the plasma membrane in root epidermal cells of Col-0, *loki044* and *det3* (G). Asterisks indicate statistical significance between mutant lines and the WT, \*\*\**P* < 0.001. *P*-values were calculated using a Student’s t-test. Representative images of FM4-64 signal at 1h and 4h in all genotypes are shown in (F). Scale bar = 100µm, arrowheads indicate tonoplast staining.

The inhibition of TOR is also known to induce changes in cell wall and cell wall integrity maintenance^22^. Notably, rapamycin treatment of ScFKBP12-expressing Arabidopsis plants leads to a decrease in cell wall rhamnogalacturonan-I content, one of the main components of pectin^6^. Pectins influence cell wall rigidity but also cell-to-cell adhesion. We used ruthenium red staining to investigate cell adhesion defects in *loki* mutants. Some loss of cell adhesion could be seen on the hypocotyls of seedlings grown in control conditions (Fig. S12A) but these defects were strongly worsened in AZD-grown seedlings (Fig. S12A, Fig. 6E). Ruthenium red staining was similarly increased upon TOR inhibition in the *det3* mutants (Fig. S12A). Consistently, cell wall content in uronic acids, mostly represented by galacturonic acid, a major component of pectic polysaccharides, was decreased in *loki* mutants on AZD (Fig. 6C). Interestingly, while metabolic profiling showed numerous changes in sugars upon AZD treatment in *loki*, cell wall composition in neutral sugars was only slightly affected by TOR inhibition (Fig. S12B). Together, these results show an additive effect of the *loki* mutation and TOR inhibition which results in defects in both cell wall rigidity and cell-to-cell adhesion.

## Discussion

The initial goal of this study was to identify new components of the plant TOR signaling pathway using a pharmacogenetics approach. Indeed, specific TOR inhibitors such as AZD have been developed in the animal field and have been shown to be active in plants^16^. These molecules have been useful to identify new actors of the TOR pathway like the YAK1 kinase that we also identified thanks to a suppressor screen^17,18^. Our first screen was performed on LY to try to identify mutants both related to TOR and PI3K kinases. The initial mutant we isolated had a peculiar phenotype of ectopic cell clusters which also appeared when plants were treated with the TOR-specific inhibitor AZD. The mutation responsible for this phenotype was named *loki* and mapped as a nonsense mutation in the coding sequence of a plant homolog of the central component of the yeast and animal RAVE complex that regulates V-ATPase activity.

In yeast and animal cells, activity of the V-ATPases can be dynamically regulated in response to nutrient levels or energization state of the cell by a conserved reversible dissociation mechanism^57^. Signals like glucose or amino acid deprivation induce the dissociation between V_1_, which is released into the cytosol, and V ^58,59^. The subunit C (VHA-C) also detaches from the rest of V ^58^. This dissociation blocks V-ATPase activity. The holoenzyme can then be rapidly reassembled to restore V-ATPase activity. This reassembly is mediated by the RAVE complex in yeast^39,60^ and by its homologue, the Rabconnectin-3 complex, in animals^61^. In yeast, the RAVE complex is composed of three subunits: Rav1, Rav2 and Skp1^45^. In animals, the Rabconnectin-3 (Rbcn3) complex was first characterized as a heterodimer consisting of Rbcn3A/DMXL and Rbcn3B/WDR7^62^, with recent studies identifying the human ROGDI protein as a third subunit of the complex^63,64^. A hybrid RAVE complex comprised of Rav1 and a WDR7 ortholog, Wdr1, was also identified in fungi^65^. Although these complexes differ in size and organization, they all have in common the presence of a Rav1 domain-containing protein at their core. Moreover, in those organisms, RAVE-mediated reassembly between V_1_ and V_0_ involves the formation of a complex between RAVE subunits and free cytosolic VHA-C^67^.

In plants, reversible dissociation of the V-ATPase has never been observed *in vivo*. However, destabilization of the peripheral stalk, a prerequisite for reversible dissociation in yeast, has been observed in Arabidopsis protoplast^66^ and in *Kalanchoë daigremontiana*^67^. This destabilization is associated with a decreased V-ATPase activity and, interestingly, involves an important conformational change of VHA-C. This plant mechanism could correspond to an evolutionary ancestral mechanism from which evolved reversible dissociation in yeast and animals.

We have identified, in Arabidopsis, the LOKI protein which shares homology with Rav1 and DMXL/Rbcn-3A, especially at the Rav1 domain. Yeast *rav1*Δ knockouts display acidification defects similar to those seen in V-ATPase mutants^45^. Here, we showed that *loki* knockouts have acidification defects associated with a decreased V-ATPase activity. These defects resemble those of the V-ATPase *det3* mutant, which carries a weak allele reducing VHA-C levels^40^. This suggests the existence of a RAVE-like complex in plants, with LOKI as the central member, possibly involved in the dynamic regulation of V-ATPase activity .

Our results also show the *loki* mutation to affect the regulation of V-ATPase activity specifically at the TGN/EE. Compartment-specificity of RAVE-mediated V-ATPase activity regulation has been described in yeast. Indeed, the VHA-a subunit Stv1-containing complexes, that localize to the Golgi and the endosomes, do not dissociate upon glucose starvation, unlike vacuolar Vph1-containing V-ATPases^30,31^. The plant RAVE-like complex could thus be compartment-dependent as is the case in yeast, with, unlike other known RAVE complexes, a specificity to the TGN/EE. The divergent evolution of this compartment-specificity between yeast and plant could be explained by the differences in the organization and functions of their endomembrane systems. Indeed, the plant TGN/EE is a one-of-a-kind compartment among eukaryotes. It has been described as a hybrid compartment, with newly-synthetized proteins, recycled proteins going to and coming from the plasma membrane, and proteins destined to the multi-vesicular bodies (MVBs) for degradation, passing through it^68^. As such, it is the main hub for protein trafficking in plant cells. Given this major role, the RAVE-like regulation of V-ATPases in plant cells could have evolved into a specific regulator of TGN/EE integrity.

One cannot exclude that the Arabidopsis RAVE complex is also affecting the vacuolar V-ATPase. However, a specificity of plant vacuoles is their bearing two distinct classes of proton pumps, the V-ATPases and the vacuolar H^+^-PPases, which are absent in yeast and animals and are also involved in vacuolar pH regulation^69^. This redundancy of the proton pumping function at the vacuole could also explain the apparent lack of effect of the LOKI loss-of-function and *det3* mutation at this compartment.

In plants and animals, V-ATPases at the TGN/EE are known not only to be involved in compartment acidification, but also in directly regulating vesicular trafficking by interacting with small GTPases and by regulating the biogenesis of organelles like MVBs^70,71^. On top of the pH defects, we also identified in *loki* mutants signs of a delayed endocytic trafficking, supporting our hypothesis of the specific role of this protein in TGN/EE regulation. Interestingly, *loki* displayed only a slight reduction of etiolated hypocotyl growth in the dark which contrasts with the de-etiolated phenotype of *det3* with short hypocotyls^40^. The de-etiolated phenotype of *det3* has been attributed to a brassinosteroid (BR) insensitivity resulting from improper recycling of the BR receptor BRI1, itself due to the delayed intracellular trafficking^41^. This could suggest a lesser impact of the *loki* mutation on endocytic trafficking or an involvement of LOKI at specific subpopulations of TGN/EE vesicles. More studies would be needed to dissect the precise effect of the *loki* loss-of-function on intracellular vesicle movement. Nevertheless, hypocotyl growth in the dark is markedly inhibited by TOR inhibition with AZD when compared to the control plants (Fig. 3). TOR inhibition has been shown to restrict hypocotyl growth and this result supports the hypothesis that *loki* mutants are hypersensitive to TOR inhibition.

Accordingly, TOR inhibition leads to a worsening of leaf growth retardation and acidification defects in those mutants, as well as a hyper-accumulation of stress metabolites (Fig. 3-4). Although interplay between TORC1 and RAVE-mediated V-ATPase regulation has been described in mammals^25^ and in yeast^28^, the link between TOR and V-ATPases in plants remains poorly understood but several connections could already be made. For instance, it is known in Arabidopsis that (FAB1)/FYVE finger-containing phosphoinositide kinase produce phosphatidylinositol 3,5-bisphosphate (PI(3,5)P_2_) which are important for the maturation of late endosomes^72^. In yeast it has been shown that TOR can phosphorylate FAB1 which causes it to shift to signaling endosomes where it generates PI(3,5)P_2_ that triggers their maturation^73^. We could then hypothesize that a combination of V-ATPase and endosome maturation defects linked to alterations in, respectively, the RAVE complex and TOR activities, could synergistically reduce trafficking at the TGN/EE.

In yeast, TORC1 signaling involves phosphorylation of the protein kinase Sch9 (a homologue of the plant S6 kinase) at the vacuolar membrane^27^. Sch9 serves as a main mediator of anabolism during TOR activation and notably interacts with V_1_ in the presence of glucose. Upon glucose depletion, Sch9 dissociates from V_1_ and this dissociation seems linked with a decrease in V-ATPase assembly^28^. These data suggest the existence of a negative feedback loop between RAVE-mediated V-ATPase reversible assembly and nutrient sensing by TORC1. In animals, interconnectedness between TORC1 and V-ATPase activity has been more precisely characterized. In mouse embryonic fibroblasts (MEFs), mTORC1 inhibition was shown to promote the activity of the RAVE complex and thereby promote the assembly of V_1_ domains from a cytosolic pool with inactive V_0_ domains. As a result, treatment with a TOR inhibitor decreases lysosomal pH^25^.

Here, our results suggest that AZD treatment has no effect on vacuolar pH in Arabidopsis. As our measurements were performed on seedlings grown for several days in the presence of the inhibitor, compensatory mechanisms cannot be ruled. We also showed that AZD treatment leads to a worsening of the TGN/EE acidification defects in *loki*. In MEFs, loss of WDR7 lead to an increase in lysosomal pH under both control and Torin1 treatment conditions^74^. However, mTORC1 inhibition does not cause a further pH increase in *wdr7* KO MEFs^74^. More data are needed as studies suggest that V-ATPase regulation by nutritional and metabolic cues can happen in both directions depending on the time-frame of the experiment^75^.

Looking at the response of V-ATPase mutants to TOR inhibition, we found that *loki* and *det3* shared the ectopic cell cluster phenotype, whereas the vacuole-specific double mutant *vha-a2xvha-a3* did not. This furthers our hypothesis of a specific interplay between the TOR pathway and V-ATPases at the TGN/EE.

In yeast, inhibition of TORC1 inhibits the activity of the plasma membrane H+-ATPase (PM-ATPase) Pma1, involved in the export of proton from the cytosol. Conversely, TORC1 activates Pma1 and disruption of this regulation leads to an acidification of cytosolic pH^76^. Similarly to the feedback loops described between V-ATPases and TORC1, Pma1 was also shown to be required for TOR activation^77^. Pma1 orthologues have been identified in tobacco^78^ and recent studies have shown a similar activation of TORC1 by stimulated PM-ATPases in *Nicotiana tabacum* BY-2 cells^79^. Given this, the inhibition of TOR in *loki* and *det3* mutants could lead to an inhibition of PM-ATPases, resulting in an accumulation of protons in the cytosol. In control conditions, PM-ATPases are functional, excess protons are exported toward the apoplast to maintain intracellular pH. However, upon TOR inhibition, those PM-ATPases could be inhibited as seen in yeast, leading to growth defects and possibly osmotic imbalances. This osmotic imbalance, in association with weakened cell walls, could be the mechanism behind the ectopic cell cluster phenotype.

Indeed, we also show that, similar to *det3* mutants^41^, *loki* display changes in cell wall composition, including a decreased cellulose content (Fig. 6), worsened upon TOR inhibition like the reduction of hypocotyl growth in the dark. We propose this to stem from a decreased trafficking of cellulose synthase cesA complexes with additive defects from the disrupted TOR pathway. Indeed, studies converge toward a central role of the TOR complex in the maintenance of cell wall integrity (CWI)^22^. Consistently, the transcriptomic profile of *loki* mutants show an enrichment in cell wall biogenesis associated genes but also suggest an over-expression of genes involved in response to wounding. Moreover, *loki* mutants display defects in cell adhesion, again worsened by TOR inhibition (Fig. 6), which are reminiscent of the defects observed in the pectin-deficient *quasimodo1* and *2* mutants^80^. Interestingly, the *quasimodo2* (also known as *tumorous shoot development 2*) mutant produces disorganized tumor-like growth^81^.

Similarly, cell cluster-like structures were observed in Col-0 seedlings recovering from heat shock and characterized as a wound response mechanism^47^. We thus propose that the development of those ectopic cell clusters in *loki* and *det3* could be the result of a combination between pre-existing defects in those mutants (like pH of the TGN/EE vesicles linked to reduced BR signaling), the loss of CWI maintenance and a dysregulated wounding response. The preferential localization of ectopic clusters on leaves and more specifically on vascular tissue remains to be explained and further studies are needed to fully understand the biological processes underlying the onset of this phenotype in the *loki* line.

## Materials and Methods

### Plant material and growth conditions

All *Arabidopsis thaliana* mutants and transgenic lines used in this study are in the Columbia genetic background, unless otherwise indicated. The following lines were used: the T-DNA mutants *loki044*, *loki109* and *loki129* (SALK_044233, SALK_109847 and SALK_129764c respectively), and the previously described *lst8-1* mutant^46^, *det3* mutant^40^ and *vha-a2xvha-a3* double mutant^38^. For TGN/EE pH measurements, *loki044* and *loki129* mutants were crossed with the SYP61-pHusion reporter line^41^. Seedlings were grown on Arabidopsis medium (2.5mM KH_2_PO_4_, 2mM MgSO_4_, 1mM CaCl_2_, 0.05% 2−(N-Morpholino) ethanesulfonic acid sodium salt (MES, w/v), 1X microelements^46^, 0.25mM Iron Ammonium Citrate) containing – unless otherwise indicated – 5mM KNO_3_, 0.8% agar (w/v), 1% sucrose (w/v) and adjusted to pH 5.7 with KOH. Seeds were sterilized in a solution of Bayrochlore and ethanol 96° for 5-10min, washed twice in ethanol 96° and left to air-dry under a sterile hood. Seeds sowed on agar plates were stratified for 1 or 2 days at 4°C to synchronize germination and then transferred to controlled growth chambers (22°C, 16h/8h light-dark cycle) for different periods of time depending on the experiment. For hypocotyl etiolation experiments, vertical plates are placed under light for 3h30 then moved to a dark chamber.

### TOR activity measurements

For TOR activity measurements, 5-day-old seedlings were grown on solid Arabidopsis medium and then transferred for 24h starvation in liquid Arabidopsis media without sucrose. Seedlings were then transferred for 5h to medium supplemented with 1% sucrose (w/v) with or without 1μM AZD. The level of RPS6 phosphorylation was used as a read-out of TOR activity as described in ref. 82. Detailed procedure for these experiments is provided in *SI Appendix*.

### Metabolomic and transcriptomic analyses

Metabolite levels were determined by global analysis after derivatization and injection into a GC-MS. *In vitro* grown seedlings were harvested 6 or 24 days after sowing on Arabidopsis medium (+/- 0.25µM AZD). Four to five biological replicates were analyzed for each mutant and control lines. Details of sample preparations, metabolite quantifications, and analyses have been previously published^18^. Raw metabolomic data are provided in Dataset S1.

Transcriptomic analyses were performed using RNA-seq and RNA extracted from three independent biological replicates. Total RNA was extracted from 18-day-old seedlings grown on solid Arabidopsis medium (+/- 0.25µM AZD) Details of RNA quantification and statistical analyses have been previously published^18^. Data from this experiment were added to the CATdb database according to the international standard MINSEQE ‘minimum information about a high-throughput sequencing experiment’ and transmitted to the NCBI database Geomnibus. For micro-dissected samples further details regarding RNA samples preparation and sequencing are provided in *SI Appendix*.

#### *In vivo* pH measurements

For vacuolar pH measurements, Arabidopsis seedlings were grown for 6 days on Arabidopsis medium (5mM KNO_3_, 1% sucrose, pH 5.7). Seedlings were incubated in liquid Arabidopsis medium containing 10μM BCECF,AM [(2’,7’-Bis-(2-carboxyethyl)-5(6)-carboxyfluorescein],(acetoxymethylester) (Molecular Probes, Invitrogen) in the presence of 0.02% Pluronic F-127 (Molecular Probes, P3000MP) for 1h in the dark at room temperature, and washed twice for 5min with Arabidopsis medium. Imaging was done on a Leica SP8 using a HCX PL APO CS 20x/0.70 IMM UV objective. BCECF was excited sequentially at 458nm and 488nm. Emission was detected between 510 and 550nm. All of the images were exclusively recorded within the elongation zone of the root. Images were processed with ImageJ and ratio values were calculated using the RatioloJ plug-in^83^, the intensity was measured for each pixel and the values of the 488nm-excited image were divided by the values of the 458nm-excited image. The ratio was then used to calculate the pH on the basis of a calibration curve.

For pH measurements in the TGN/EE, SYP61-pHusion seedlings were grown for 7 days on Arabidopsis medium (5mM KNO_3_, 1% sucrose, pH 5.7). Imaging was done on a Leica SP8 using a HC PL APO CS2 63x/1.20 water objective. The pHusion construct was excited at 488nm and 561nm. GFP fluorescence was detected between 510 and 540nm, RFP fluorescence was detected between 600 and 670nm. All of the images were recorded within the elongation zone of the root. Images were processed with ImageJ, signal was first segmented for TGN/EE using the automatic ‘Moments’ threshold, then ratio values were calculated using the RatioloJ plug-in. The intensity was measured for each pixel and the values of the 488nm-excited image were divided by the values of the 561nm-excited image. Calibration procedures for BCECF and the pHusion probe are provided in *SI Appendix*.

### V-ATPase activity measurements

V-ATPase activity of microsomal membranes was colorimetrically determined by following Pi release as previously described^38^ with minor modifications. Microsomal fractions were prepared from 7-day old seedlings grown on Arabidopsis medium (5mM KNO_3_, 1% sucrose, pH 5.7). Plant material was ground in liquid nitrogen and resuspended in homogenization buffer (350mM sucrose, 70mM Tris-HCl (pH 8.0), 3mM Na_2_EDTA, 0.15% (w/v) BSA, 1.5% (w/v) PVP-40, 4mM DTT, 10% (v/v) glycerol and 1X protease inhibitor mixture (c0mplete mini, EDTA-free, Roche). The homogenate was filtered through two layers of Miracloth (Calbiochem) and centrifuged at 15 000*g* for 15min at 4°C. The supernatant was then centrifuged at 100 000*g* for 1h. The resulting microsomal pellet was resuspended in resuspension buffer (175mM sucrose, 10mM MOPS-Tris (pH 7.0), 2mM DTT and 1X protease inhibitor mixture). V-ATPase activity of 10μg microsomal membranes was determined as phosphate (Pi) release after 30min incubation at 28°C. The V-ATPase assay solution contained 25mM MOPS-Tris (pH 7.0), 4mM MgSO_4_, 50mM KCl, 1mM NaN_3_, 100μM Na_2_MoO_4_, 1mM Na_3_VO_4_, 0.02% Brij-35 and 3mM Mg-ATP. V-ATPase activity was calculated as the value difference between the measurements in the absence and presence of 200nM Concanamycin A plus 50mM KNO_3_. Reactions were terminated by adding 40mM citric acid. Freshly prepared AAM solution (50% (v/v) acetone, 2.5mM ammonium molybdate, 1.25M H_2_SO_4_) was then added to the reaction and absorbance was immediately measured at 355nm. 10μg of BSA instead of fresh microsomes were used for the blank value.

### Cell wall biochemical composition analyses

Cell wall composition was assayed on hypocotyls of 14-day-old seedlings dark-grown on Arabidopsis medium with or without 0.25µM AZD. The monosaccharide composition of the non-cellulosic fraction was determined by hydrolysis of alcohol insoluble residue (AIR) with 2M trifluoroacetic acid (TFA) for 1h at 100°C and the crystalline cellulose fraction was obtained after H_2_SO_4_ hydrolysis of the TFA-insoluble fraction. The monosaccharides of both fractions were quantified by HPAEC PAD on a Dionex ICS-5000 instrument (Thermo Fisher Scientific) equipped with a CarboPac PA20 analytical anion exchange column (3mm x 150mm). Detailed procedure for cell wall samples preparation and analysis is provided in *SI Appendix*.

### Cell wall confocal imaging

For cell wall staining and cluster imaging, aerial parts of seedlings were incubated overnight at room temperature in fixation solution (75% ethanol (v/v), 25% acetic acid (v/v)) under gentle shaking. Fixed samples were washed twice in PBS 1X (13.7mM NaCl, 0.27mM KCl, 1mM Na_2_HPO_4_, 0.2mM KH_2_PO_4_), cleared overnight in ClearSee solution^84^ and then stained in the dark in ClearSee solution supplemented with 0.1% Calcofluor or 0.1% Direct Red 23. Before imaging, samples were washed three times for 30min in PBS 1X. For 2D cluster imaging (Direct Red 23), leaves were mounted in water and imaged using a Leica SP5 confocal microscope with an HC PL APO 20x/0.75 water objective. For whole cluster 3D imaging (Calcofluor), leaves were fixed to the bottom of a petri dish with solidified agar 4% (w/v), covered in water and imaged using a Leica SP8 confocal microscope with an HC FLUOTAR L 25x/1.0 IMM long working distance objective. Images were acquired as Z-stacks and processed through the ImageJ Z-project tool.

### Statistical analyses

Unless otherwise stated, all experiments were repeated at least twice. Statistical analyses were performed using either two-tailed Student’s t-test (P < 0.05), Wilcoxon rank test (P < 0.05) or Dunn test (P < 0.05) adjusted with BH correction on Rstudio. Details of calculated *P*-values, associated statistical outputs and number of samples are indicated in the legends of the *Figures*. Data visualization was performed with the Rstudio packages ggplot2 (version 3.5.1) and tidyplots (0.2.1).

Detailed procedures for phenotypical parameters measurements, RT-PCR, microdissection, FM4-64 uptake analysis, Ruthenium Red staining and imaging, and PI/FDA dual staining are provided in *SI Appendix*.

## Acknowledgments

We would like to thank Pr. Karin Schumacher and Dr. Melanie Krebs (COS, Heidelberg, Germany) for providing the Col-0 / SYP61-pHusion and *det3* / SYP61-pHusion lines, and for their assistance in setting up V-ATPase activity assays. Additionally, we thank members of the OV-Cytology/Imaging platform (IJPB, Versailles, France) for their help with the various imaging experiments.

S.L. and C.I. are supported by Ph.D. grants from the Fondation de la Recherche Médicale (FRM ECO202306017358 and ECO201806006346). J.B. was supported by the ANRT (Association Nationale Recherche Technologie, #2022/1139) and the Fertinagro France company. C.F., C.M. and A.-S.L. were partly funded by the DecoraTOR ANR grant (ANR14-CE19-007). This work is partly supported by a French State grant (Saclay Plant Sciences, ANR-17-EUR-0007, EUR SPS-GSR) managed by the French National Research Agency under an Investment for the Future program (reference n◦ ANR-11-IDEX-0003-02).

